# Ensembles for improved detection of invasive breast cancer in histological images

**DOI:** 10.1101/2023.04.13.536542

**Authors:** Leslie Solorzano, Stephanie Robertson, Johan Hartman, Mattias Rantalainen

## Abstract

Accurate detection of invasive breast cancer (IC) can provide decision support to pathologists as well as improve downstream computational analyses, where detection of IC is a first step. Tissue containing IC is characterized by the presence of specific morphological features, which can be learned by convolutional neural networks (CNN). Here, we compare the use of a single CNN model versus an ensemble of several base models with the same CNN architecture, and we evaluate prediction performance as well as variability across ensemble based model predictions.

Two in-house datasets comprising 587 WSI are used to train an ensemble of ten InceptionV3 models whose consensus is used to determine the presence of IC. A novel visualization strategy was developed to communicate ensemble agreement spatially. Performance was evaluated in an internal test set with 118 WSIs, and in an additional external dataset (TCGA breast cancer) with 157 WSI.

We observed that the ensemble-based strategy outperformed the single CNN-model alternative with respect to accuracy on tile level in 89% of all WSIs in the test set. The overall accuracy was 0.92 (DICE coefficient, 0.90) for the ensemble model, and 0.85 (DICE coefficient, 0.83) for the single CNN alternative in the internal test set. For TCGA the ensemble outperformed the single CNN in 96.8% of the WSI, with an accuracy of 0.87 (DICE coefficient 0.89), the single model provides an accuracy of 0.75 (DICE coefficient 0.78)

The results suggest that an ensemble-based modeling strategy for breast cancer invasive cancer detection consistently outperforms the conventional single model alternative. Furthermore, visualization of the ensemble agreement and confusion areas provide direct visual interpretation of the results. High performing cancer detection can provide decision support in the routine pathology setting as well as facilitate downstream computational analyses.

## Introduction

Breast cancer is one of the most commonly diagnosed cancer diseases [1]. The high incidence of breast cancer (BC) cases poses a substantial burden on healthcare providers, including pathologists. To alleviate some of the clinical burden, decision support systems based on computer vision and machine learning can be utilised [2], including systems designed for characterization and diagnosis of BC from routine digital histopathology whole slide images (WSI) from tissue sections stained with Haematoxylin-eosin (H&E).

Accurate detection and coarse segmentation of invasive cancer (IC) in breast histology is of special importance since the IC component is central in several downstream tasks, including e.g. histological grading [3]. IC detection also enables delineating of regions of interest in whole slide images (WSIs), and detection of the IC component also often serves as the first step for other modeling tasks, where downstream analyses are performed only in the IC region in the H&E image[4]. However, digital H&E WSIs of resected tissue sections are of very large sizes and high resolution (100k x 100k pixels; 0.25 μm per pixel), thus manual annotations and comprehensive visual inspection are both challenging and time-consuming. At the same time, there is also a well known high interobserver variability in histopathology assessments [5], which cause a degree of uncertainty in general, and can in the worst case lead to the wrong diagnosis and possibly cause suboptimal choice of treatment [6].

Due to these practical challenges there is a need to develop machine learning (ML) and deep learning (DL) methodologies that can be applied for analyses of H&E images to assist with the detection and segmentation of breast cancer in WSIs.

Deep CNN models have previously been developed for breast cancer detection in the context of the BreaKHis data set [7] (N=82 breast tumours) [8–10]. For further previous work in the domain, please see recent review articles [11,12].

Most of the previously reported studies focusing on breast cancer detection are based on very small datasets (less than 100 WSIs) and single models. However, single models have intrinsic uncertainty in both their parameters and the predictions, which are associated with the optimisation strategy as well as intrinsic uncertainty in the data used to train the models. In situations where individual models are associated with a degree of uncertainty, a well-known strategy to reduce variance in predictions is to use ensemble learning across a set of base models [13][14]. Several studies in the area of computational pathology employ ensemble learning [15][16][4] to improve prediction performance. Using an ensemble model can also provide a measure of agreement between base models.

In this study, we propose an ensemble-based modeling approach to improve breast cancer cancer detection and coarse segmentation, and we systematically compare performance between a single deep CNN modeling approach for IC detection [4], and an ensemble learning alternative. The study is based on training and test data sets that are substantially larger than previously reported studies, thus providing more conclusive results.

## Methods

### Data and preprocessing

For our experiment we used two in-house datasets, Clinseq-BC [17] and SöS-BC, a retrospective cohort of patients diagnosed at the Stockholm South General Hospital (Sweden) [4]. From Clinseq, 232 (185 training, 47 test) WSI with manual digital annotations were included, and from SöS 355 (284 training, 71 test) WSI with manual digital annotations were included. This translates to a total of 2,502,649 tissue tiles of 598×598 pixels (271×271 μm) at 20X magnification. Validation was performed in the independent external dataset TCGA-BC [18], 157 WSIs with digital annotations by [19], including 1,589,216 tiles.

Tiles in the in-house datasets were preprocessed as described previously in [4]. Both datasets include WSI level annotations of routine clinicopathological factors, including the associated Nottingham Histological Grade (NHG) and routine IHC biomarker status for ER, HER2 and Ki67 proteins. The Clinseq dataset contains exhaustive spatial annotations of IC specifically, and the SöS dataset contains exhaustive annotations for IC and Ductal Carcinoma In-Situ (DCIS) and partial annotations for other types of BC and artifacts. Such annotations are overlapped and a single tile can contain several labels which have to be mapped to IC vs the rest. For TCGA-BC the IC annotations were obtained from the work in [19].

### Ensemble learning

There are three main aspects to ensemble learning, (1) data sampling/selection; (2) training each base-model of the ensemble; and (3) combining results (Polikar 2012). We explored different combinations of the CNN base-models of the ensemble.

### Sampling and selection for diversity

The available WSIs from both in-house datasets were separated into 80% training and 20% is held out as an internal test dataset. The training data was then divided into 10 equal groups, keeping the proportion of clinical features. Each base model is trained with one of such divisions. These divisions make it so that each base-model in the ensemble is optimised to classify IC and not-IC, finding different features and making different decisions.

### CNN as base-models of the ensemble

Following the experiments performed in [20] where Inception V3, ResNet50, Inception-ResNet V2 and Xception models were studied. Inception V3 yielded the best results in terms of classification of H&E tiles of prostate cancer biopsies. Therefore Inception V3 was deemed suitable to provide a separation of histological patterns. Each network is trained starting with ImageNet weights and then optimized with the training data until convergence or early stopping decided by the tuning set. The loss function is cross entropy and the AdaBelief [21] optimizer was used. The default parameters for Inception V3 were used as implemented in PyTorch.

A decision about whether a tile contains IC or not depends on the output probability for each CNN in the ensemble. Although the output is continuous, as it is normalized with a softmax function, we cannot guarantee that the CNNs are calibrated. For this reason, the CNN output is converted to a binary label but not with 0.5 as threshold. To select an appropriate threshold, first: all CNN in the ensemble are applied to all the training data. Second: for each CNN a threshold is applied which optimizes the dice score for all the data divisions in which said CNN was not trained, e.g CNN 1 is trained with division 1 and after training, the calibration is performed with its predictions on divisions 2-10.

### Combining CNN results

When binarized, the outputs of the CNN can be seen as a vote. In such case, the chosen aggregation was majority voting. As shown in [22] for a number of classifiers *T*, each with a probability of correct classification *p*, the accuracy of the ensemble follows a binomial distribution. The probability of ensemble success is eq 1, where *k* >=*T*/*2*+*1*

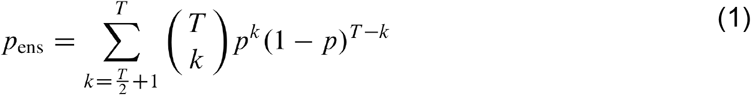

*p*_*ens*_ approaches 1 as *T* → ∞if each individual *T* has a probability of success *p* higher than 0.5.

### Using Ensemble predictions as a measure of agreement

The prediction by each base-model CNN can be considered as a “vote” for a certain class, and the level or agreement, or variance, can be seen as a measure of uncertainty on a decision. A high level of base-model agreement indicates that several base-models agree despite that they were trained on different subsets of the data. This measure can be used as a metric to guide interpretation of the prediction results.

We define agreement as the amount of base-models that “vote” for the correct class. If threshold *t*_*i*_ is the binarization of the *ŷ*_*i*_ output of base-model *f*_*i*_ using threshold *τ* and the correct label is *l*, then the agreement *a* of *n* base-models over a tile *x* can be written as:

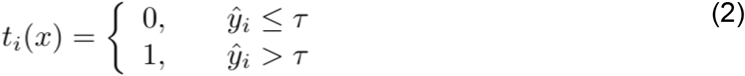

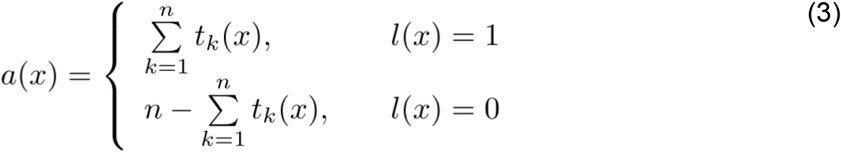

In this way, agreement will always be a number between *0* and *n*, where *0* denotes complete disagreement with the ground truth and *n* will be the maximum number to represent complete agreement with the ground truth.

## Results

To assess if an ensemble modeling strategy can improve breast cancer detection in routine WSIs stained with hematoxylin and eosin, we implemented an ensemble modeling strategy and compared it with the single model equivalent.

Individually, every base-model CNN has a high probability of correct classification. Moreover, the base-models are trained on different data, and have intrinsic variability between them. Therefore, from the theory we can expect that an ensemble will provide higher accuracy, which was also observed here. In Figure 1. the plots show the accuracy comparison in each internal test set. Similarly, Figure S1 shows a similar trend for dice score across WSI.

**Figure 1.**
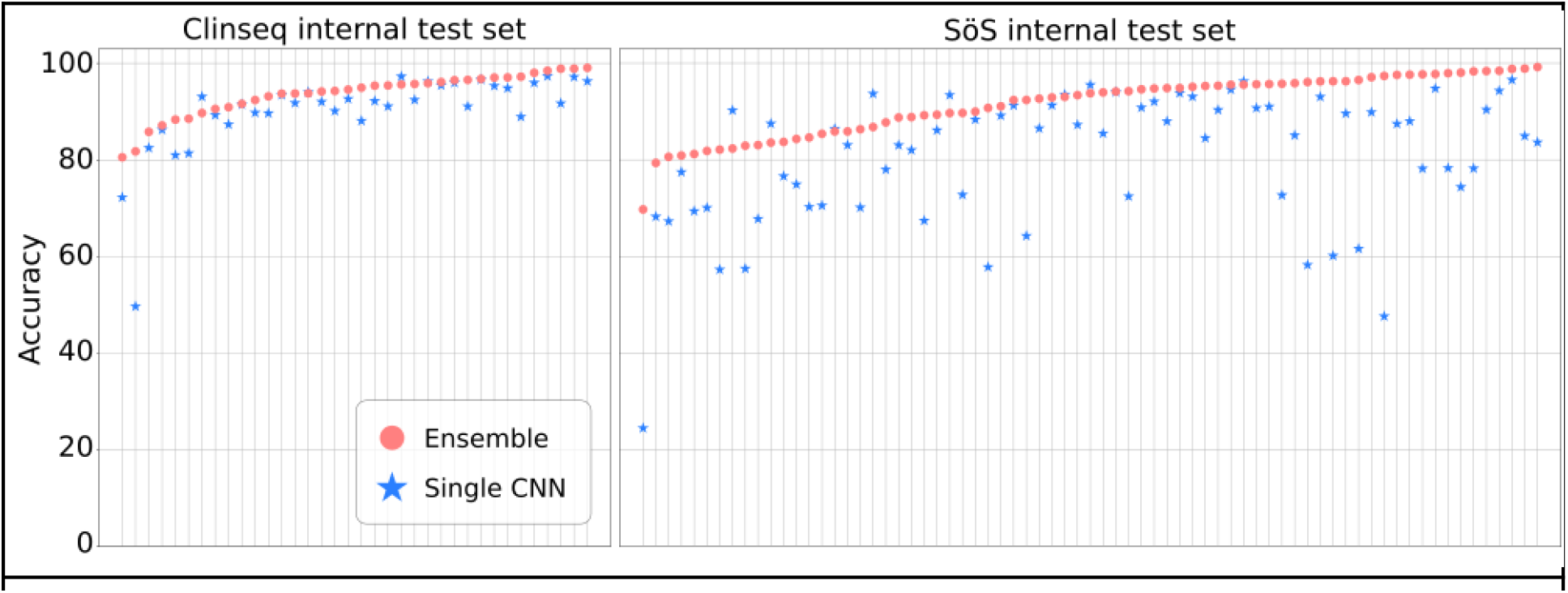
Accuracy in internal test sets. Pink markers show the accuracy achieved by the ensemble in the 2 internal test sets, for each WSI. Blue markers show the results of the single network. At a glance the large number of blue markers under the pink area shows that the ensemble achieved a higher proportion of accuracies per WSI.

First, performance was assessed in the internal test set, we found that ensemble models consistently outperformed the single-model alternative (Table 1). With a CNN ensemble we achieved 92.97% (2,326,937 / 2,502,649 tiles) accuracy across both in-house test datasets. In Clinseq (internal test set) the ensemble model provided better accuracy compared to the single CNN model alternative in 88.8% of WSIs, and in SöS-BC (internal test set) 88.7%.

**Table 1.** Summary statistics of ensemble results vs single model results in test data

|  | Ensemble SöS | Single model SöS | Ensemble Clinseq | Single model Clinseq | Ensemble TCGA | Single model TCGA |
| --- | --- | --- | --- | --- | --- | --- |
| Accuracy | 0.91215 | 0.804946 | 0.935704 | 0.900575 | 0.874758 | 0.753852 |
| Dice Score | 0.861954 | 0.764373 | 0.946297 | 0.915296 | 0.892073 | 0.780212 |
| Specificity | 0.858872 | 0.609806 | 0.852501 | 0.930852 | 0.927529 | 0.442272 |
| Precision | 0.836572 | 0.677470 | 0.958906 | 0.977182 | 0.829357 | *Not available* |

Next, we evaluated the generalizability of our proposed ensemble model in an external and fully independent data set (TCGA breast cancer), and we confirmed that the ensemble performance was consistently higher than the single model alternative in 96.8% of the WSIs. Figure 2 shows the performance comparison in the external dataset. We also assess performance using dice score (Figure S2), confirming a similar trend.

**Figure 2.**
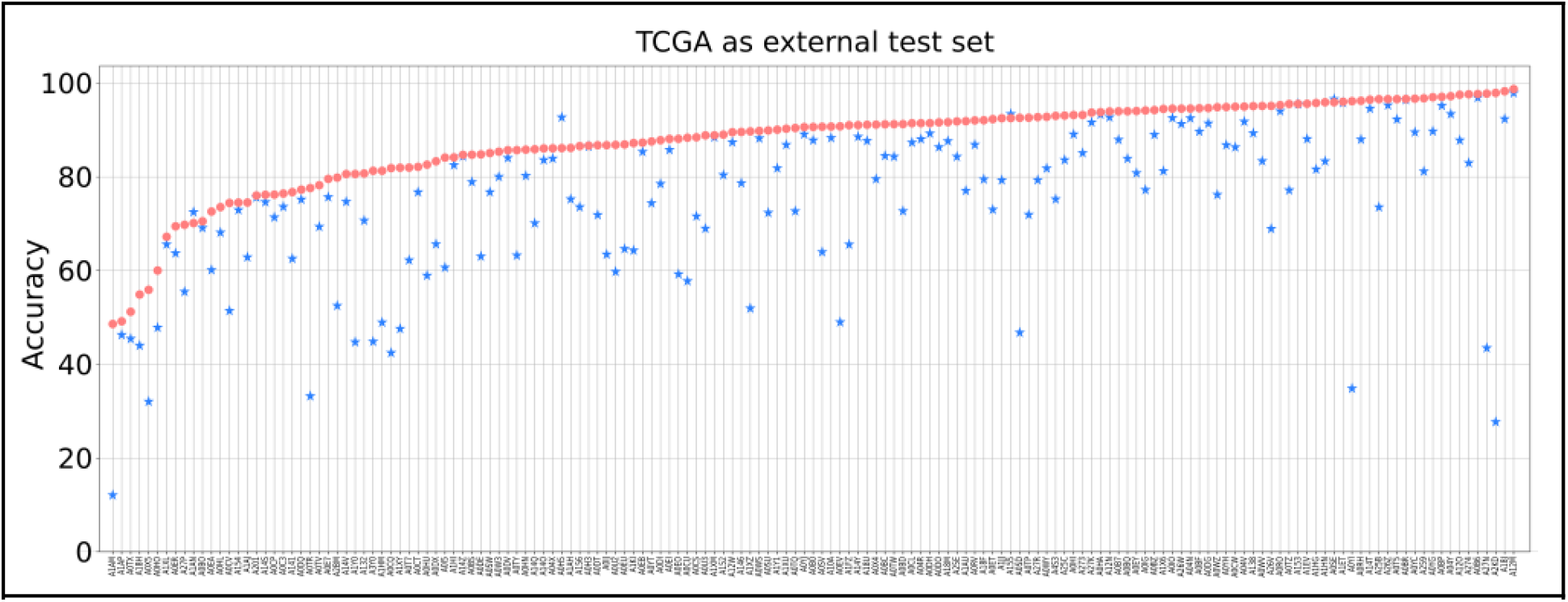
Accuracy in the external test set. Pink markers show the accuracy achieved by the ensemble in the TCGA dataset, for each WSI. Blue markers show the results of the single network. At a glance the large number of blue markers under the pink area shows that the ensemble achieved a higher proportion of accuracies per WSI.

### Interpretation and visualisation of ensemble prediction results

To explore how the agreement in ensemble predictions could aid interpretation of prediction results, we developed a novel visualisation strategy. Figure 3 shows the agreement plotted along with the ground truth and the results of the ensemble. It allows for an immediate view of locations where there was disagreement and a measure of how much of it there was. Additionally three example tiles are shown, on B there is a texture that is clearly different from C and D. B is correctly classified with high agreement. C actually presents a tile labeled as IC but the ensemble predicted incorrectly and with many CNNs voting for the wrong class (hence the dark color for agreement). In D is a tile correctly classified as not IC with high agreement. However C and D look similar to the untrained eye, and there is a chance that simply the CNNs didn’t see or didn’t learn features that allowed for the separation of the two types of tiles.

**Figure 3.**
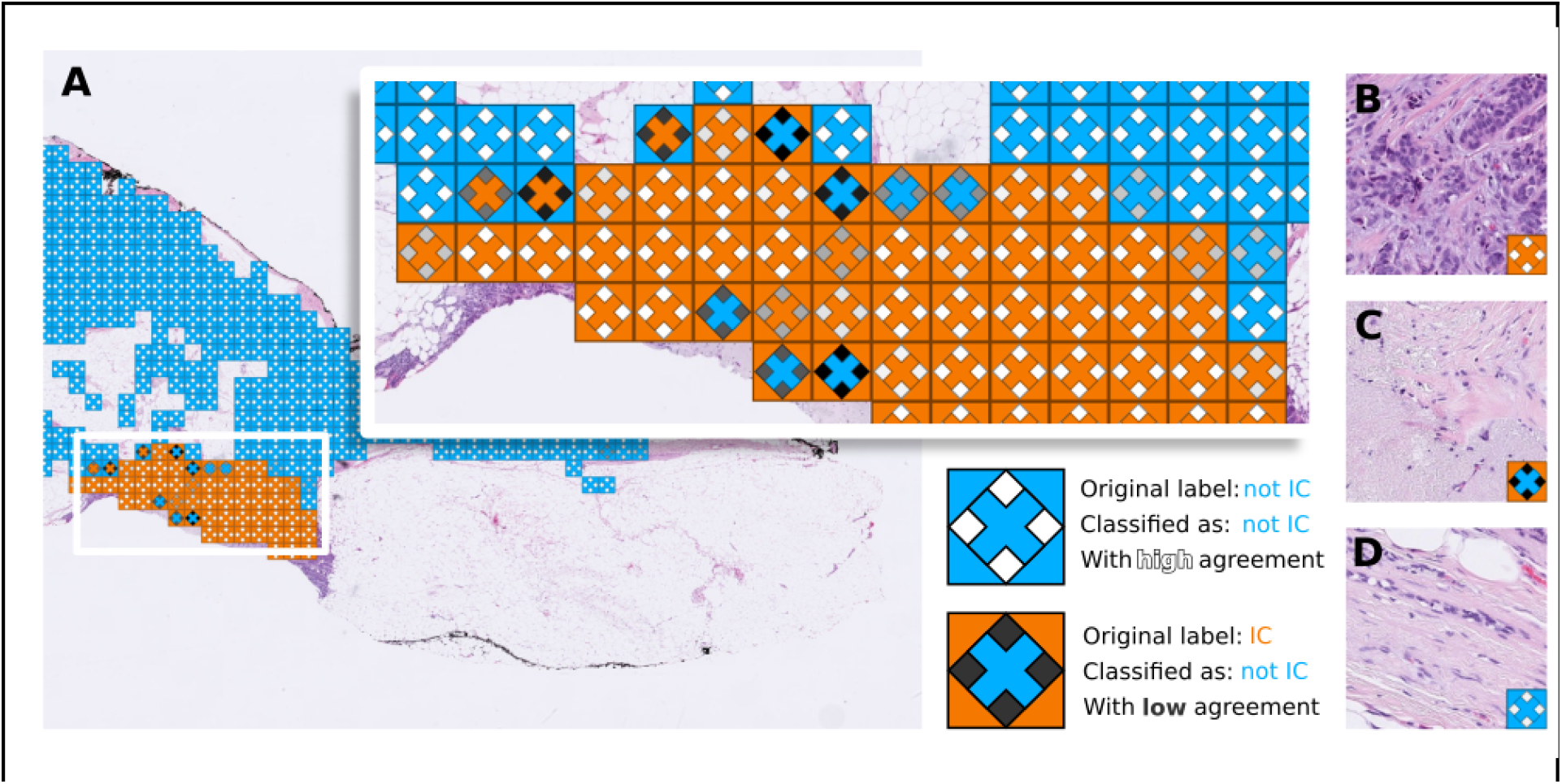
Spatial visualisation of ensemble results and base-model agreement. A.) Overlaid in the WSI are the original label as a square, a diamond representing agreement mapped in greyscale where white is agreement 10, which means 10 networks voted for the right class. And an X with the predicted label. Orange means IC and blue means not IC. Example tiles include B.) a high agreement in a tile with IC, C.) a tile that is labeled as IC but the prediction is to be not IC. D.) a tile without IC. This visualization allows for a quick examination of tiles that have contradictory labels or disagreements.

Next, we aimed to explore potential benefits of the ensemble modelling strategy. Figure 4 shows an exploration of the behavior of agreement and the confusion within the labels SöS dataset, DCIS is of particular interest. Figure 4A. shows that 10 out of 10 networks voted for the correct class in a large portion of the tiles, which means that 10 out of 10 CNNs were able to learn different patterns that allow for a good separation of IC vs non-IC. We expected a confusion between IC and DCIS due to their differences being not in the morphology but in their location and surroundings, however the large imbalance in the amount of tiles representing the classes makes it difficult to draw conclusive insight.

**Fig 4.**
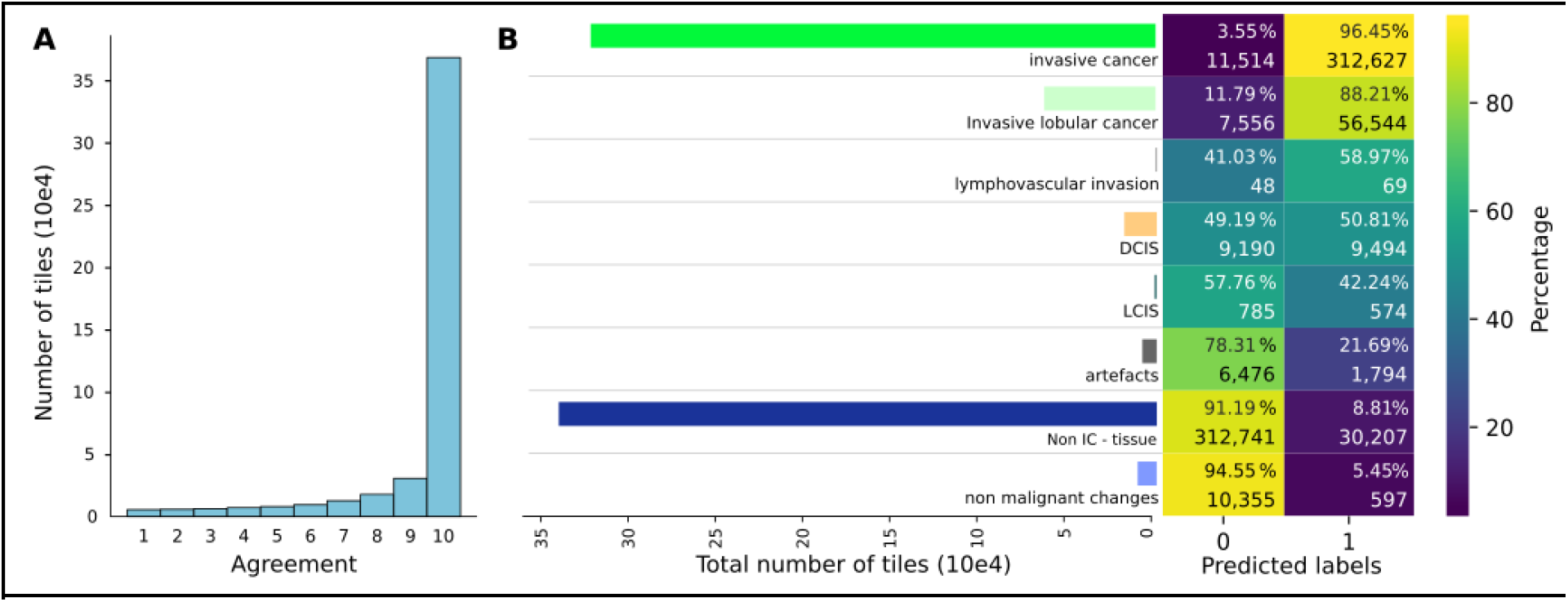
Agreement. A) The histogram of agreement amongst test set tiles. Shows that there is a high level of agreement regardless of the class. B) Confusion matrix between SöS classes and IC/non-IC label. Only the labels “invasive cancer” and “invasive lobular cancer” should have high amounts of IC. The confusion matrix’s colors represent the percentage, which is also written, along with the number of tiles it represents. Additionally the horizontal bar chart represents the total amount of tiles available for each of the labels.

## Discussion

In this article we developed and benchmarked a deep learning-based ensemble modeling strategy for for the detection of IC in breast cancer WSI. Our goal was to take advantage of the properties of ensembles, particularly for binary classification, which enables us not only to improve the accuracy and dice score but also to explore a measure of agreement of the base-models of the ensemble and relate it to a certainty on the predicted label which can be taken into account for further downstream analysis. Additionally we present a brief exploration into the relationship of IC with other labels that were available in one of our datasets, namely DCIS.

In terms of sample size, we believe that the 587 WSI that were available in the present study in the internal datasets might be a limitation. However, compared to most previous studies based on e.g. the BreakHis dataset, the present study is still >7 times larger with respect to patients (WSIs), and over 400 times larger with respect to the number of tiles with exhaustive annotations of IC and not-IC.

It should be noted that this work is done on a per tile basis which is relevant to most applications [11]. Originally the annotations come in the form of polygons (series of coordinates within the WSI) that can be overlaid on the image, and the labels per tile are given based on the position of the center of the tile within such polygons. Even when a minimum of tile area is expected to be inside the polygon annotation, the actual amount of IC in it can be a source of noise, and even original labels could be explored in terms of the agreement in them, particularly in the borders of an IC region.

Referring back to three main aspects to ensemble learning according to (Polikar 2012), ((1) data sampling/selection; (2) training each base-model of the ensemble; (3) combining results) (1) we split the training data such that the distribution of clinical features NHG grade, ER, HER2 and Ki67 were kept proportional in an attempt to reduce any variability from these factors. (2) The choice of CNN model is not a trivial task, as there are not only a plethora of models available but they are all capable of capturing a wide range of high-dimensional features to separate IC from the rest. Several models have been tried in histology for different classification tasks but following the experiments performed in [20] where Inception V3, ResNet50, Inception-ResNet V2 and Xception models were studied. Inception V3 yielded the best results in terms of classification of H&E tiles of prostate cancer biopsies. Therefore Inception V3 was deemed suitable to provide a separation of histological patterns. However there remains a possibility of exploring different architectures for the base-models. (3) The outputs of each CNN base-model are continuous and to obtain a decision as an ensemble they can be aggregated as the mean or median and then thresholded. However, binarizing before aggregation presented the benefit of a majority voting strategy, not only allowing us to visualize the decision but also to support the decision with the framework in equation 1 for binary distributions, where so long as every base-model has a chance higher than 0.5 of success then there can be a higher probability of obtaining the right classification for each tile in the WSI.

Although visually interesting, agreement is not a direct explainability measurement. It does not explain the reasons why a decision is made, but it instead provides a particular type of metric relating to the agreement between a set of CNN models, which can be interpreted as the concordance in learned representations across the CNNs in the ensemble.

Detection of IC remains an important task in routine pathology, and computer-based models have the potential to provide decision support to pathologists in this work. Detection of IC is also a first step for downstream analyses and modeling of histopathology WSIs, where it is common that the main interest is in the analysis of the invasive cancer component. In this study we systematically compared and evaluated the performance between single CNN models and an ensemble alternative. We observed strong support for the application of an ensemble approach. In future work it would be interesting to also investigate explainability further, and also systematically evaluate the impact of the IC detection step on other downstream analysis tasks.

