## Supplementary figures for "Ensembles for improved detection of invasive breast cancer in histological images": Suppl of Ensembles for improved detection of invasive breast cancer in histological images.pdf

Leslie Solorzano, Stephanie Robertson[2], Johan Hartman[2], Mattias Rantalainen[1]
[1]Department of Medical Epidemiology and Biostatistics, Karolinska Institutet, Stockholm, Sweden
[2]Department of Oncology-Pathology, Karolinska Institutet, Stockholm, Sweden

### Supplementary material

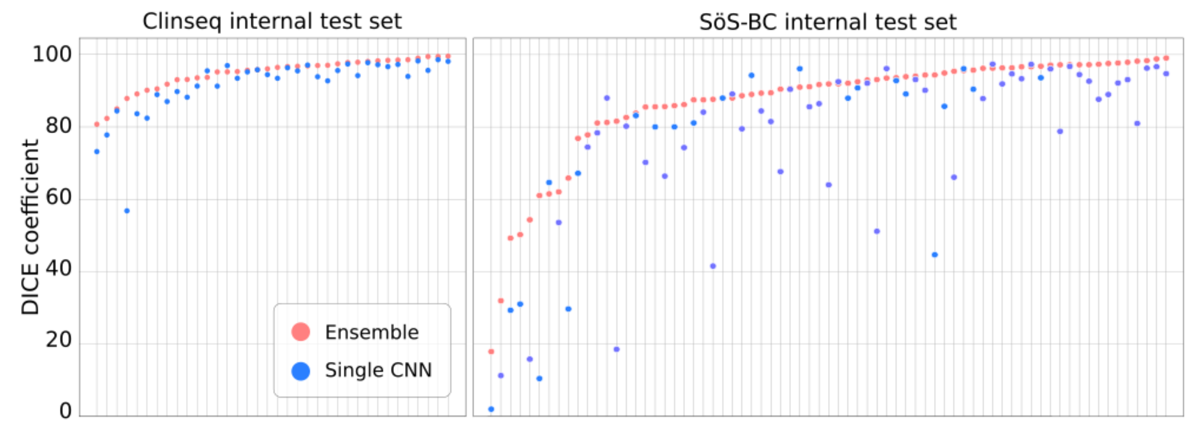


Figure S1. **Dice coefficients in internal test sets.** Pink markers show the dice coefficient achieved by the ensemble in the internal test sets for each WSI. Blue markers show the results of the single network. At a glance the large number of blue markers under the pink area shows that the ensemble achieved a higher proportion of accuracies per WSI.

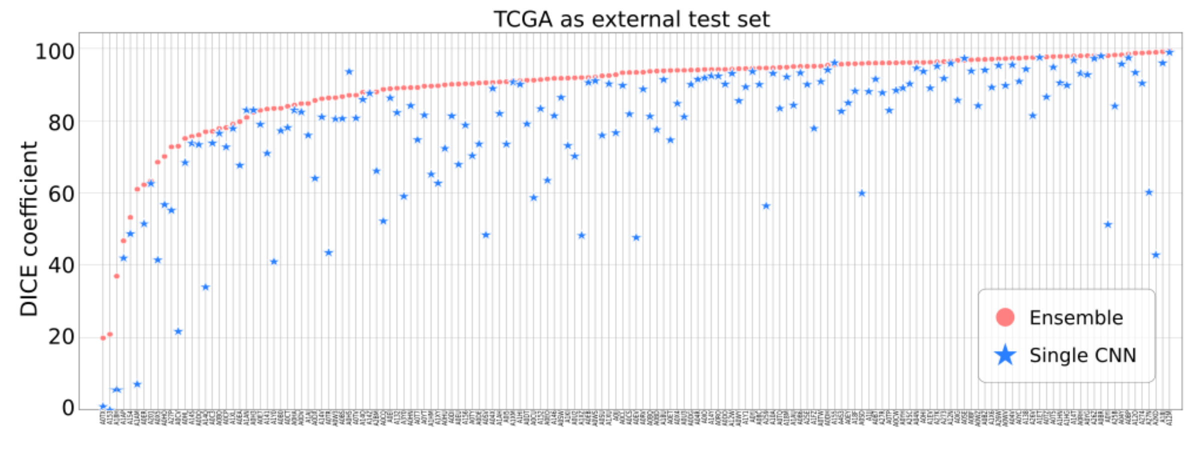


Figure S2. **Dice coefficients in the external test set.** Pink markers show the dice coefficient achieved by the ensemble in the TCGA dataset, for each WSI. Blue markers show the results of the single network. At a glance the large number of blue markers under the pink area shows that the ensemble achieved a higher proportion of dice coefficients per WSI.

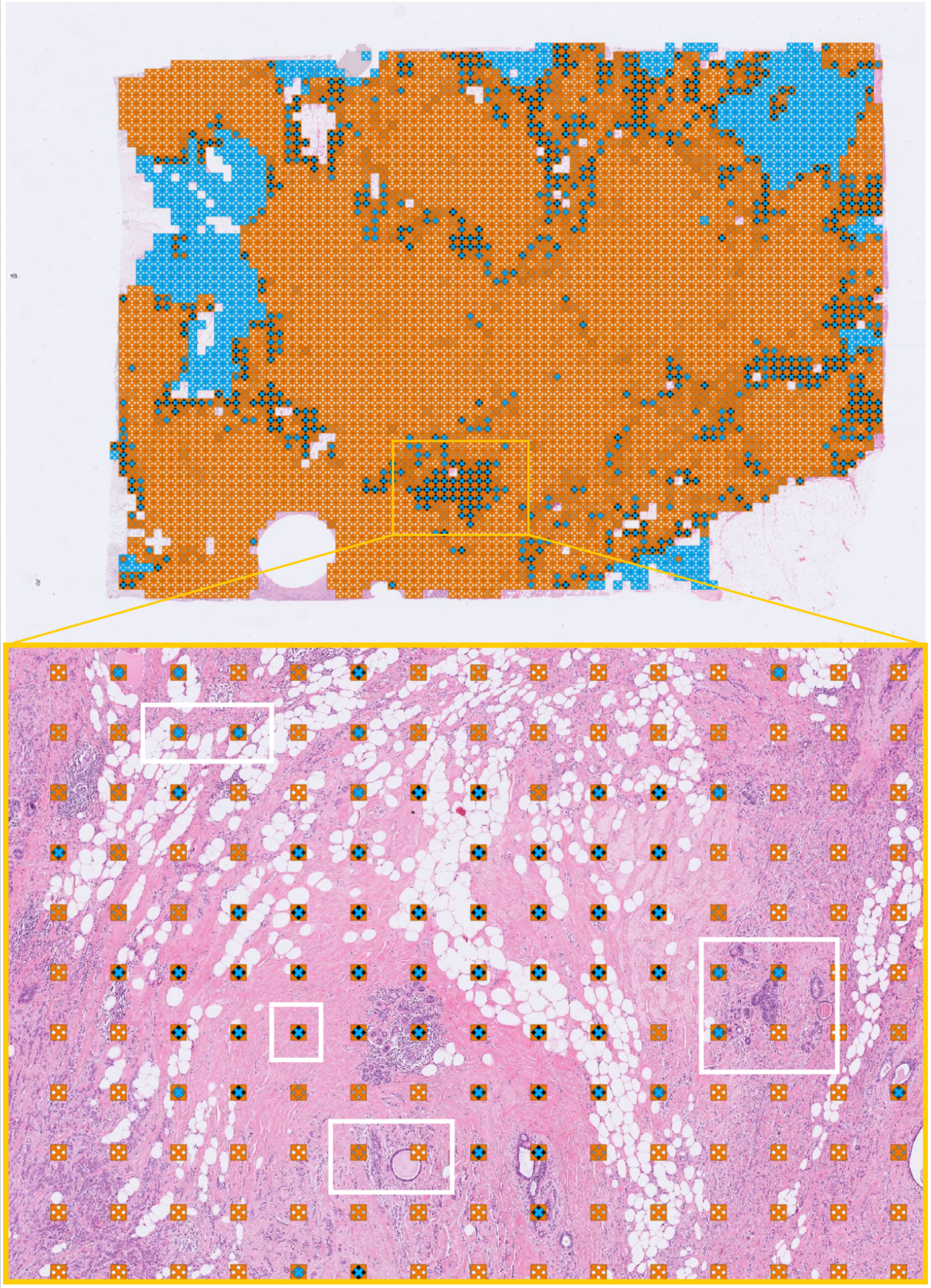

Figure S3. Agreement on WSI where a single model surpassed the ensemble accuracy. We want to explore why this could be. Note the agreement of lower than 10 (gray colors) along the border of a big IC region (orange). An enlarged area is highlighted and the tissue underneath is observed, note that there is adipose tissue present and structures marked in white.

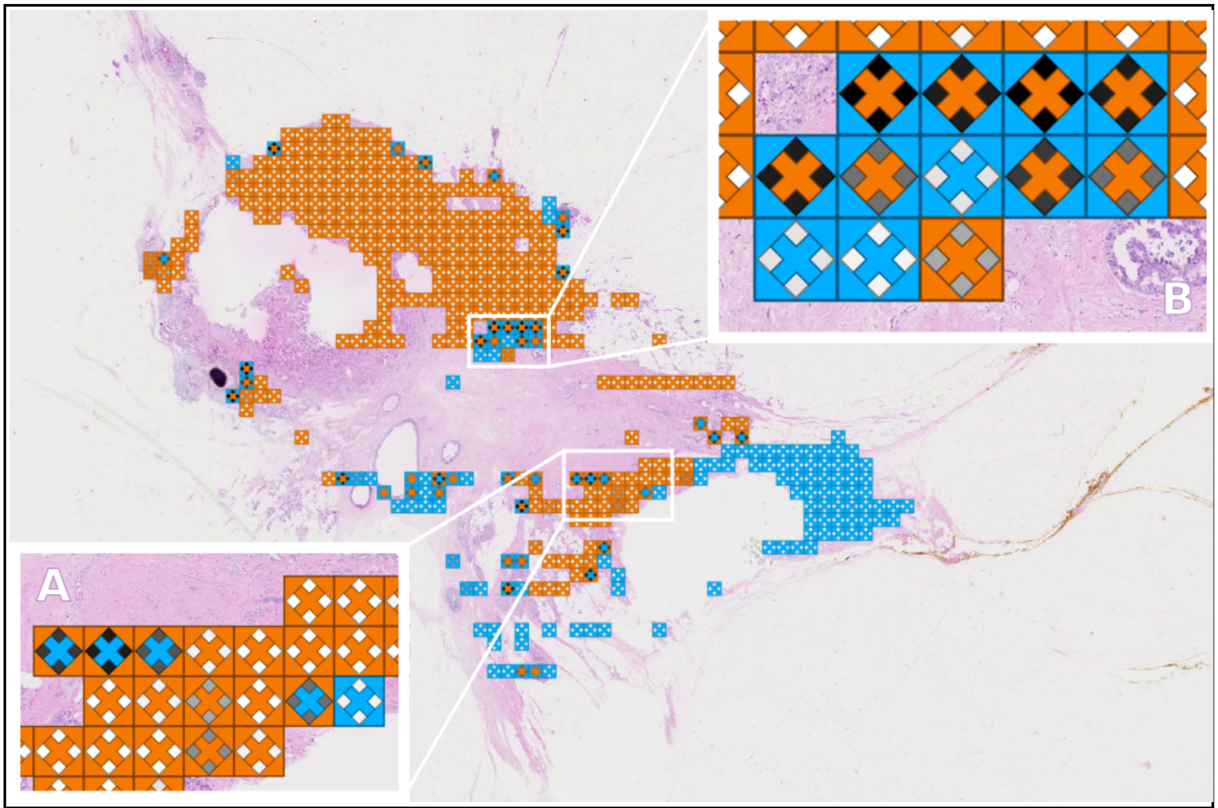


Figure S4. Visualization of agreement of a WSI where the difference in accuracy is small: 87.5% vs 83.56%. In A) a few tiles contain IC as ground truth while the predicted value is non-IC, with a gray agreement meaning it disagreed but not extremely. There are also some correctly labeled IC where not all CNN voted for IC. In B) a strong disagreement between labels indicates that the CNNs vote for IC, however the ground truth indicates non-IC. This kind of tile can be reviewed by humans.
